# Heat pre-treatment reduces multiplicity of plasmid transformations in yeast during electroporation, without diminishing the transformation efficiency

**DOI:** 10.1101/2024.07.03.601847

**Authors:** Marcus Wäneskog, Emma Elise Hoch-Schneider, Shilpa Garg, Christian Kronborg Cantalapiedra, Elena Schaefer, Michael Krogh Jensen, Emil Damgaard Jensen

**Affiliations:** The Novo Nordisk Foundation Center for Biosustainability, Technical University of Denmark, DK-2800 Kgs. Lyngby, Denmark

**Keywords:** DNA transformation, DNA electroporation, Automated genotyping, heat-shock, barcodes, yeast, plasmids, DNA libraries

## Abstract

High-throughput DNA transformation techniques are invaluable when creating high-diversity mutant libraries, and the success rate of any protein engineering endeavors is directly dependent on both the size and diversity of the mutant library that is to be screened. It is also widely accepted that in both bacteria and yeast there is an inverse correlation between the DNA transformation efficiency and the likelihood of transforming multiple DNA molecules into each cell. However, most successful high-throughput mutant screening efforts require high quality libraries, i.e., libraries comprised of cells with a clear phenotype-to-genotype relationship (one genotype/cell). Thus, DNA transformation methods with a high multiplicity of transformation are highly undesirable and detrimental to most mutant screening assays. Here we describe a simple, robust, and highly efficient yeast plasmid DNA transformation methodology, using a dual heat-shock and electroporation approach (HEEL) that generates more than 2 x 10^7^ plasmid-transformed yeast cells per reaction, while simultaneously increasing the fraction of mono-transformed cells from 20% to more than 70% of the transformed population. By also using an automated yeast genotyping workflow coupled with a dual-barcoding approach, consisting of a SNP and high-diversity barcode (10N), we can consistently identify and enumerate unique plasmid genotypes within a heterogeneous population merely through Sanger sequencing. We demonstrate here that the size and quality of a transformed library no longer need to be inversely correlated when transforming large mutant DNA libraries in yeast using highly efficient DNA electroporation methods.

**Significance:** With the recent expansion of artificial intelligence in the field of synthetic biology there has never been a greater need for high-quality data and reliable measurements of phenotype-to-genotype relationships. However, one major obstacle to creating accurate computer-based models is the current abundance of low-quality phenotypic measurements originating from numerous high-throughput, but low-resolution assays. Rather than increasing the quantity of measurements, new studies should aim to generate as accurate measurements as possible. The HEEL methodology presented here aims to address this issue by minimizing the problem of multi-plasmid uptake during high-throughput yeast DNA transformations, which leads to the creation of heterogeneous cellular genotypes. HEEL should enable highly accurate phenotype-to-genotype measurements going forward, which could be used to construct better computer-based models.

## Introduction

Numerous studies have investigated the molecular mechanisms that affect DNA transformations in both bacteria and yeast. In the case of chemical transformation of bacteria, the primary parameters of importance are temperature (heat-shock), the presence of certain metal ions (Rubidium, Manganese and Calcium) and/or pH, as reviewed in (1). Most studies conclude that chemical transformation induces either a semi-permeabilized and/or transiently permeable membrane structure, or a lowering of the membrane potential, which facilitates a passive DNA uptake (1). Temperature (heat-shock) is also an important factor that stimulates extracellular DNA uptake during chemical transformation of yeast (2). However, unlike for bacteria, yeast chemical DNA transformation is currently understood to be mediated by endocytosis pathways (3), which is strongly enhanced by chemical agents, such as Lithium acetate, crowding agents (PEG) and nonspecific single-strand carrier DNA (ssDNA) (2, 3). In the case of bacterial and yeast DNA transformation mediated by electroporation, our current understanding suggests that a passive DNA diffusion through transiently formed pores in the membrane is the primary route of DNA uptake during electroporation (1, 4). Nevertheless, yeast electroporation does require a more complicated and cumbersome protocol as compared to bacterial electroporation, because the rigid yeast cell wall and/or membrane first needs to be conditioned to allow for adequate DNA uptake (5).

Clear cross-species comparisons of DNA transformations are challenging, as different factors affect the DNA uptake in bacteria and yeast. In addition, the convention to present DNA transformation outcomes as transformation efficiencies, by normalizing the number of transformed cells per µg DNA, makes direct comparisons exceptionally difficult and leads to incorrect conclusions. For bacterial DNA transformations, very small amounts of DNA are usually used, often in the range of 10-50 ng (6). Thus, when the number of transformed cells is normalized to µg of DNA, the transformation efficiency gets inflated by a factor of approximately a 100-fold. While for yeast transformations, which often use 1-10 µg of DNA, the DNA transformation efficiency becomes deflated by as much as one order of magnitude when normalized (5). Consequently, a reported transformation efficiency (CFU/µg DNA) of 10^10^ for *E. coli* means that only 10^7^ cells were transformed with 10 ng DNA per reaction. Yet, a reported transformation efficiency of 10^6^ for yeast would also mean that 10^7^ cells were transformed, but with 10 µg per reaction. Thus, transformation efficiencies are easily misunderstood and manipulated by merely changing the amount of input DNA. Also, since most studies and synthetic biology endeavors have little-to-no interest in transformation efficiencies *per se*, it is preferable and more accurate to instead report on transformation yields, i.e., the total number of transformed cells per reaction. Another important factor to consider is the fraction of transformed cells in a solution, i.e., how many cells are needed to achieve the desired transformation yield. In this study we have chosen to exclusively focus on these two parameters to best convey the true value of our results and present our findings in an easily understood and relevant format.

Transformation yields are critical when creating high-diversity mutant libraries for e.g. directed evolution purposes. However, most chemical transformation methods of either bacteria or yeast are not sufficiently high-throughput to successfully transform high-diversity libraries (1, 2). In contrast, standard electroporation protocols are often 10-100-times more efficient than most chemical transformation methods (1, 4, 5). Thus, for high-throughput random protein engineering, the transformation of high-diversity mutant libraries via electroporation is currently the most effective method. Moreover, while many studies have identified numerous important parameters that affect transformation efficacies, few studies have identified factors that control multiplicity of DNA transformations (i.e. the number of DNA molecules that enter each cell). The few studies that have investigated this phenomenon have all concluded that the likelihood of transforming a bacterial or yeast cell with multiple DNA molecules is directly correlative with the concentration of extracellular plasmid DNA that is added to the transformation reaction (6–8). This creates a challenging scenario, as transformation yields in both yeast and bacteria also increase with an increased concentration of extracellular plasmid DNA (4). Thus, with current electroporation methodologies it is not possible to design a strategy that results in both a high fraction of mono-transformed cells (good library quality), while simultaneously also retaining the maximum possible number of transformed cells (transformation yield). This inverse relationship between library size and quality is a tremendous problem for most synthetic biology approaches that rely on high-diversity mutant libraries, as the likelihood of identifying a beneficial mutant within any library is primarily limited by the size, i.e., coverage, of the library. But what is even more crucial to any library screening effort is the ability to measure and assign a clear phenotype-to-genotype relationship for each individual cell. Thus, library quality and library size are two mutually exclusive factors of equal importance. Arguably, this concern is of less importance for experiments that merely aim to identify any beneficial mutant(s), which then later can be isolated and characterized in-depth in a monoculture. However, with the increasing use of artificial intelligence (AI) and machine learning algorithms, there is now a great need for precise phenotype-to-genotype measurements to better understand the sequence space of the respective protein. The resulting high-quality datasets can then be used to train computer-based models with the aim to improve accuracy and insight of subsequent predictions (9).

In this study, we have aimed to improve the transformation efficiency of a previously published yeast electroporation protocol, while simultaneously maximizing the fraction of mono-transformed cells to achieve the largest possible high-quality mutant library of easy-to-screen yeast cells. To validate our approach, we have designed an automated yeast genotype workflow based on a single-nucleotide polymorphism (SNP) and high-diversity (10N) dual-barcoding approach that allows us to rapidly, and efficiently, identify and quantify the number of plasmids transformed in each single-cell derived yeast colony. Using this dual-barcoding approach, we discovered that subjecting yeast cells to a heat-treatment before electroporation increased the fraction of mono-transformed cells from 20% to more than 70% of the transformed yeast population. Remarkably, this heat-pretreatment increased the fraction of mono-transformed yeast cells without affecting the transformation yield, which allowed us to retain a transformation yield of up to 2 x 10^7^ yeast cells per electroporation reaction. Thus, we can overcome the classical dilemma of inversely correlated library size and library quality and maintain maximal library size while simultaneously increasing library quality, simply by introducing a heat-pretreatment.

## Results

### *S. cerevisiae* cells exhibit low survivability post-electroporation when transformed with an established DNA transformation protocol

Our initial aim was to design a barcoding approach that could be used to easily track both the unique genotype of each DNA molecule within a high-diversity mutant library and to quantify the multiplicity of transformations when transforming our model yeast, *Saccharomyces cerevisiae* (CEN.PK2-1C or CEN.PK110-10C). Toward that end, we first attempted to use a well-known yeast DNA electroporation protocol from Benatuil *et al.* to achieve ultra-high-diversity libraries in our CEN.PK2-1C yeast strain (5). To our surprise, we were not able to achieve transformation yields as reported by Benatuil *et al.* When we transformed our CEN.PK2-1C strain with 8 µg of an empty dual bacterial and yeast expression vector (pYB-Dual (10)) we observed a transformation yield of approximately 10^5^ transformed cells per reaction (Fig. 1). This is a stark contrast to the previously reported transformation yield of approximately 5 x 10^8^ transformed cells per reaction (5). However, Benatuil *et al.* transformed the common *S. cerevisiae* yeast-surface display strain EBY100 in their study, not CEN.PK2-1C. Thus, we thought that intra-strain differences might mean that CEN.PK2-1C was more difficult to transform than some other yeast strains (5). To explore this possibility, we repeated our transformation experiment with both EBY100 and the common *S. cerevisiae* strain BY4741 (Fig. 1). Surprisingly, both the BY4741 and EBY100 strains were transformed less efficiently than CEN.PK2-1C, with approximately 10^3^ and 10^4^ transformed cells per reaction, respectively. Additionally, we observed that <0.2% of EBY100, <0.6% of BY4741 and <4% of CEN.PK2-1C yeast cells survived the electroporation (Fig. 1). Benatuil *et al.* reported having 6.4 x 10^8^ starting cells per electroporation cuvette (1.6 × 10^9^ cells/mL x 0.4 mL per cuvette), of which they could transform 5.42 x 10^8^ cells per electroporation cuvette (5). Thus, they reported an ability to transform >84% of all yeast cells in solution (5). Since <4% of all *S. cerevisiae* cells, regardless of strain background, survived the electroporation procedure in our laboratory, our findings are in stark contrast to the findings reported by Benatuil *et al.* (5).

**Figure 1.**
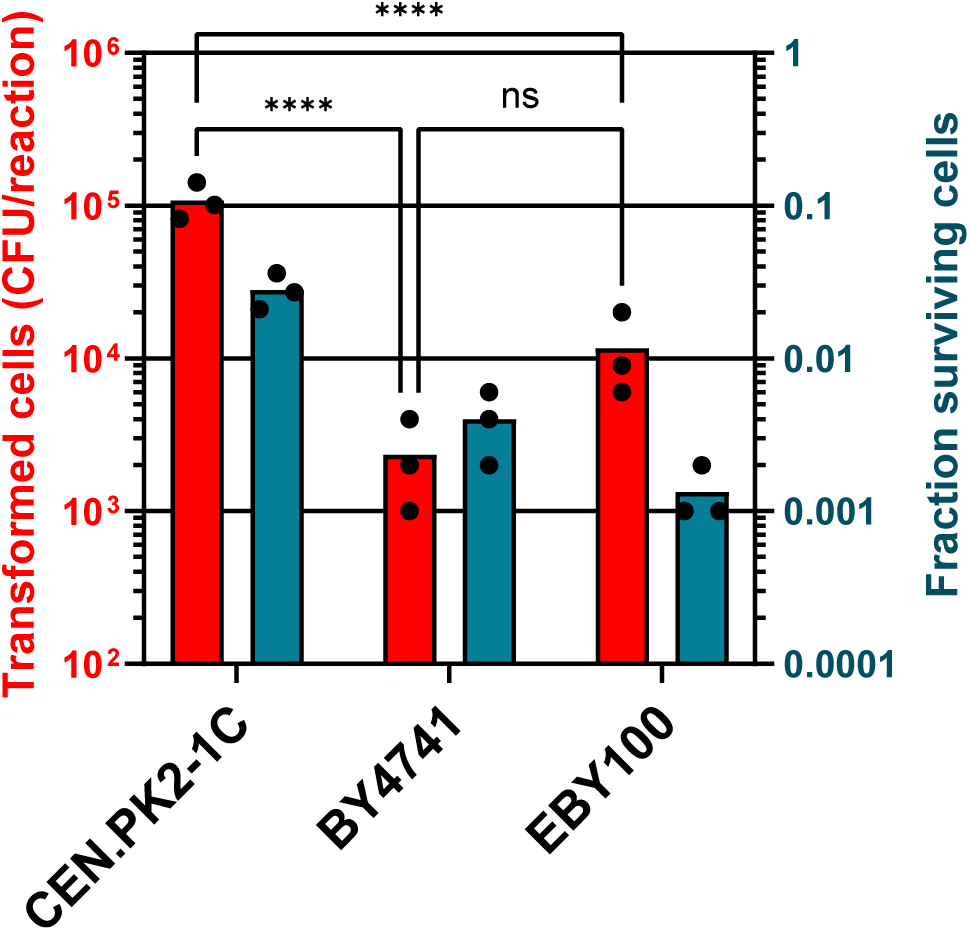
*S. cerevisiae* strains: CEN.PK2-1C, EBY100 and BY4741, have low survivability post-electroporation when transformed with a previously established protocol (5). Three different *S. cerevisiae* strains were transformed with an empty dual bacterial and yeast expression plasmid (pYB-Dual), using the method described by Benatuil *et al.* (n=3) (5, 10). Statistical significance was calculated by Two-way ANOVA with ns: P>0.05 and ****: P≤0.0001.

### Efficient *S. cerevisiae* DNA transformation by electroporation requires single-stranded carrier DNA and lower voltage

As we observed very low yeast survivability post-electroporation when attempting to reproduce the DNA transformation method described by Benatuil *et al.,* we concluded that the conditions described in the transformation protocol is too harsh on the cells (5). Following this line of thought, rather than using the EBY100 strain, we chose to continue with our preferred model strain CEN.PK110-10C. This strain is closely related to the CEN.PK2-1C strain, which displayed the greatest transformation yield (10^5^) in our lab (Fig. 1). Next, we established a permissive electroporation protocol, which we confirmed was the optimal approach, by systematically changing one parameter at a time (Fig. 2A-F). Using this optimized protocol (see Material and Methods), we could transform approximately 4% of all surviving cells in solution and achieve a transformation yield of up to 2.2 x 10^7^ cells/reaction (Fig. 3A-B). From our investigations we could confirm that both the DTT (10-20 mM) and DNA concentration (4-8 µg) previously reported by Benatuil *et al.* were also optimal in the experiments conducted in our laboratory (Fig. 2A, C-D) (5). However, we could not confirm the importance of any additional parameters previously reported (5). When we changed the electroporation voltage from 2.5 kV to 2.0 kV, almost 1-log more cells were transformed, likely because the survivability of the yeast increased from 0.05% to 6.5% of all yeast cells (Fig. 2B, Dataset 1). While an electroporation voltage of 1.5 kV did allow for even more cells to survive the treatment (12.5%), it did not increase the transformation yield compared to 2.0 kV. Thus, we observed that a greater fraction of viable cells was transformed at 2.0 kV, compared to 1.5 kV, which is a parameter of great importance for selection-free library screenings. We also observed that recovering cells after transformation in 50% sorbitol and 50% YPD did not significantly influence survivability or transformation yield (Fig. 2E) (5). Additionally, we theorized that if chemical transformation of yeast is stimulated by ssDNA, then this might also hold true for yeast when electroporated (2). As expected, the addition of 100 µg of ssDNA to the electroporation reaction increased the transformation yield >1-log when we transformed our CEN.PK110-10C strain with an empty pRS413 plasmid (Fig. 2C-D)(11). Recently, Loock *et al.* have also reported that ssDNA increases the DNA transformation yields of *S. cerevisiae* during electroporation (12). Thus, we observed that many parameters important for chemical transformation of yeast (Lithium acetate and ssDNA) are also important for electroporation of yeast. Consequently, we were curious to investigate if heat-shocking yeast cells would also affect the transformation yield during electroporation. However, when we explored the effect that a heat-shock temperature of 42°C has on yeast transformability by electroporation, we observed reduced transformation yields (Fig. 2A-D). Changing the electroporation voltage to 1.5 kV slightly compensated for this diminished transformation yield, but not to a point where we could justify a potential advantage of exposing yeast cells to a 42°C heat-shock step before electroporation (Fig. 2B). Interestingly, heat-shocking cells at 37°C before electroporation did not result in a significant difference in transformation yields, although we did observe a potential effect on the fraction of transformed cells (Fig. 2F, Dataset 1). This indicates that a mild heat shock affects the cell’s ability to take up extracellular DNA and to survive electroporation, without diminishing the overall transformation yield.

**Figure 2.**
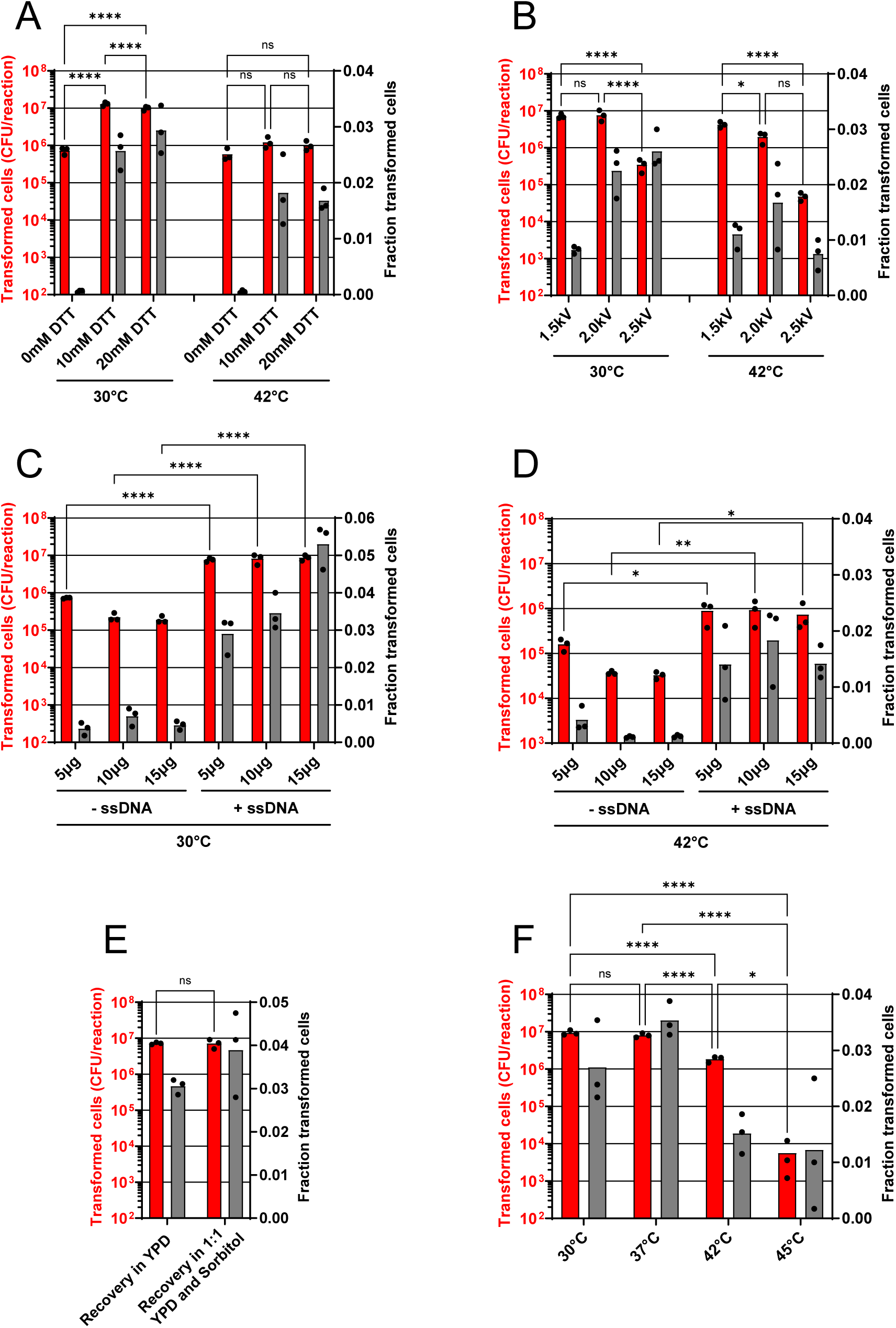
The DNA transformation method by Benatuil *et al.* can be improved by changing the electroporation voltage and by adding single-strand salmon sperm DNA (ssDNA) to each transformation reaction. *S. cerevisiae* strain CEN.PK110-10C was transformed with a pRS413 circular plasmid molecule, using our improved yeast DNA transformation method. Yeast cells were conditioned at either 30°C or 42°C, for 1h in a Lithium acetate and DTT solution (see Material and Methods). A) The effect on yeast transformability by changing the DTT concentration, B) the electroporation voltage, C-D) plasmid DNA concentration and adding ssDNA, E) omitting sorbitol from the recovery media, or F) changing the heat-shock temperature (n=3). Statistical significance was calculated by Two-way ANOVA with ns: P>0.05, *: P≤0.05, **: P≤0.005, ***: P≤0.0005 and ****: P≤0.0001.

**Figure 3.**
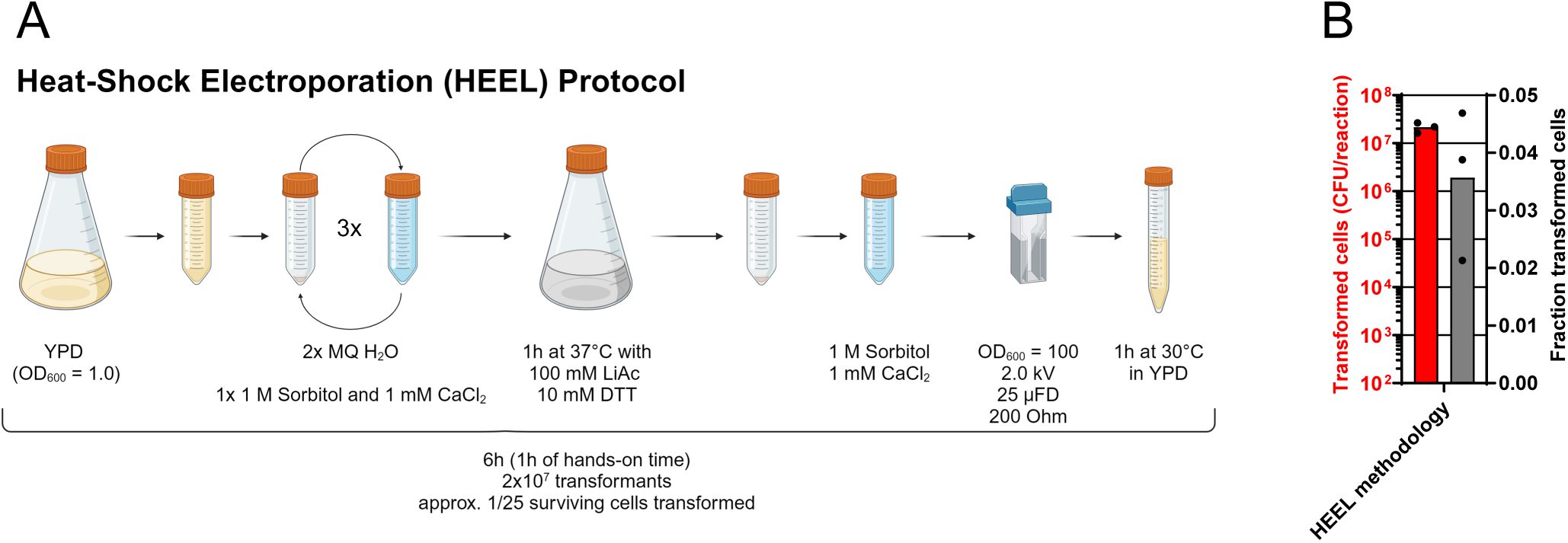
Overview of the HEEL methodology. A) Schematic illustration of the Heat-Shock Electroporation (HEEL) yeast DNA transformation method. Illustration created with BioRender.com. B) *S. cerevisiae* strain CEN.PK110-10C transformed with a pRS413 circular plasmid molecule, using the HEEL method (n=3).

### A dual-barcode approach allows for direct enumeration of plasmids post-transformation using Sanger sequencing

Multiplicity of transformation in yeast during high-throughput DNA transformations have previously been investigated (7). During this investigation, Scanlon *et al.* reported that when transformation yields were high (9.3 x 10^6^ transformed cells/reaction) then 88% of all yeast cells were transformed with multiple unique DNA molecules during electroporation (7). As our improved DNA electroporation protocol differs considerably, compared to the protocol used in that study, we were curious to investigate how our protocol affected yeast multiplicity of transformation. In their study, Scanlon *et al.* used a plasmid library consisting of a 3nt random barcode to identify how many unique plasmids were transformed into each yeast cell during electroporation (7). However, to quantify the number of unique sequences per cell, Scanlon *et al.* first performed an outgrowth of the transformed yeast cells for up to 35 hours and then identified the number of unique plasmid sequences they could find in 10-20 randomly selected and sequenced colonies. Statistical probabilities were then applied to the findings to determine the most probable number of plasmids per cell (7). This approach allowed Scanlon *et al.* to determine that most cells (88% of all transformed cells) were transformed with a median of 4 plasmids per cell (7). Drawing inspiration from Scanlon *et al.,* we designed a more comprehensive method of quantifying the multiplicity of transformation in yeast using a dual-barcode approach. This dual-barcode was designed to contain one region of 11 bp, which only contained 1 SNP per molecule, and one completely random 10 nt region (Fig. S1A). Using this dual-barcode we could directly determine the number of plasmid(s) that each yeast cell was transformed with during an electroporation by simply performing a single Sanger sequencing reaction on the region encoding the dual-barcode and then enumerating the number of unique SNPs found during sequencing. Although it should be noted that accurate SNP calling for a Sanger sequencing reaction requires that at least 15% of all sequences are identical. i.e. the maximum number of unique SNPs (plasmids) that can be identified during a single Sanger sequencing using a SNP-based method is 5-6 per cell (13). Furthermore, as no library creation method is perfect, we expected to have both empty vector contamination as well as unwanted nucleotide deletions within a few of the barcoded plasmids we created for our library. To overcome this obstacle, we designed the second 10 nt random region within the dual-barcode to act as a quality control and complementary method to the SNP base-calling approach (Fig. S1A). This 10 nt random region was designed to exploit the statistically highly unlikely event that all 10 random nucleotides were identical if multiple unique plasmids were randomly mixed (Fig. S1B). I.e., if a cell contained 2, or 3 plasmids with a random 10 nt region, then this region should contain a maximum of 2 or 3 unique sequences, respectively, at every position. While if a cell contains 4 plasmids, then at least 1 position should contain all 4 bases. The statistical likelihood of accurately quantifying the presence of 1, 2 or 3 plasmid/cell using this approach is >99%, while cells transformed with 4 plasmids will be enumerated correctly approximately 63% of times (Fig. S1B). Though this method cannot be used to determine the exact genotype of each plasmid within a library, and only has a maximum detection limit of 4 unique plasmids per single-cell-derived yeast colony, it functions well as an accurate, but low-resolution, complementary methodology to the SNP-based barcoding approach (Fig. S1B). In the case of the 11nt barcode, where only 1 SNP/molecule is present, only 33 unique sequences (11 nt x 3 possible alternative nucleotides per position) can be created (Fig. S1A). The likelihood that 2, 3, 4, 5 or 6 plasmids would have the exact same SNP is 3%, 9%, 17%, 27% and 38% respectively (Fig. S1B). Thus, while the high-diversity barcode (10N) allows for a more accurate quantification of multiplicity of plasmids in the 1-3 plasmid/cell range, the SNP-based barcode allows for a more accurate quantification in the 4-6 plasmid/cell range, as well as a precise SNP (genotype) determination (Fig. S1B). To further streamline the plasmid quantification process, we also designed an automated yeast genotyping pipeline, which allowed us to leverage the efficiency of an automated colony picker and an liquid handling robot to quickly collect, prepare, PCR amplify, and then sequence the mini-libraries found within each single-cell-derived yeast colony (Fig. 4A) (see Material and Methods).

**Figure 4.**
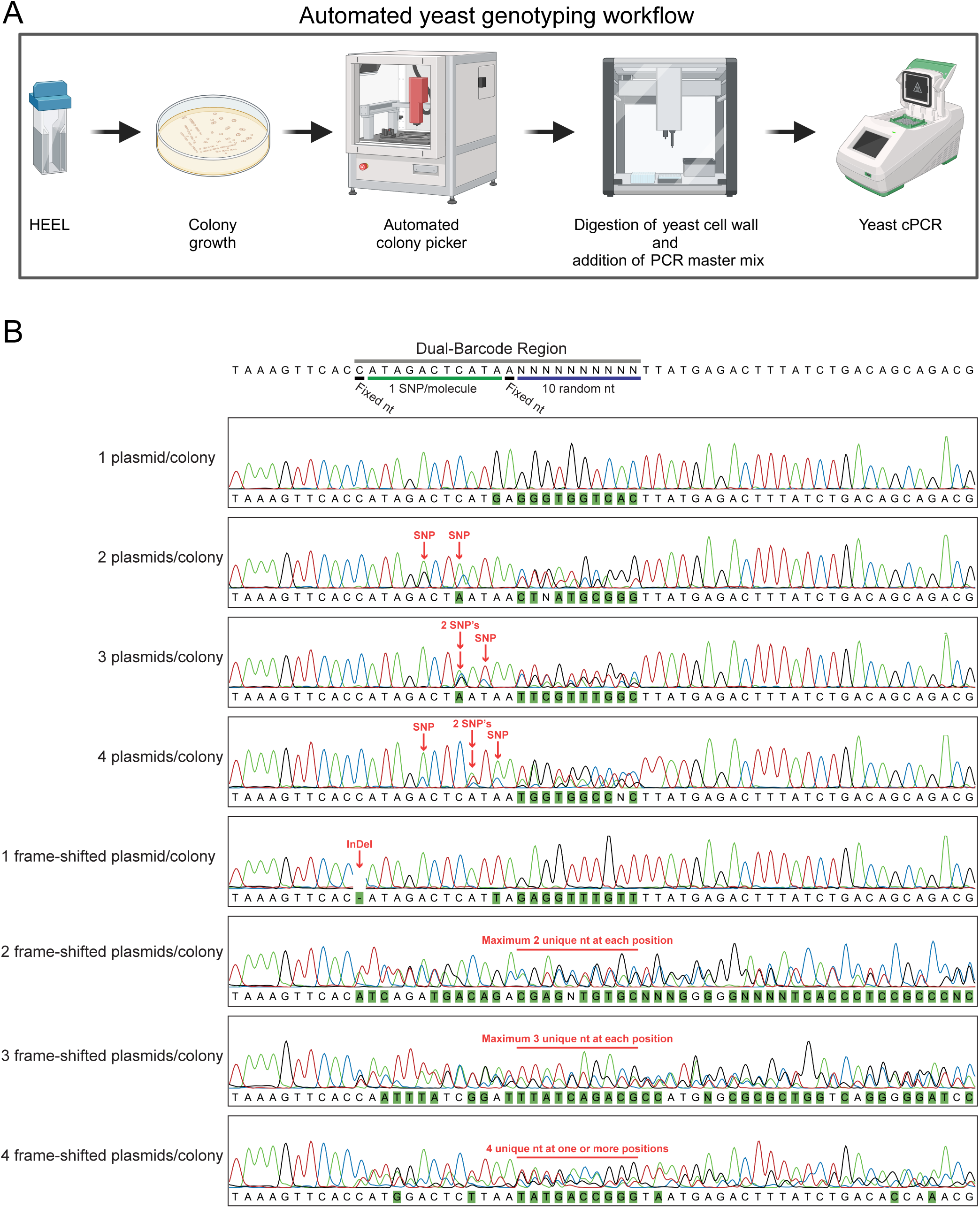
A dual-barcode approach can enumerate the number of plasmid molecules present in each single-cell-derived yeast transformant, using a single Sanger sequencing reaction. A) Schematic illustration of our automated yeast genotyping workflow. Illustration created with BioRender.com. B) Schematic illustration of the dual-barcode region and representative Sanger sequencing chromatograms from single-cell derived yeast colony PCR (cPCR). Demonstration of the utility of both the SNP and high-diversity region of the dual-barcode to enumerate the number of unique sequences in a mixed population.

Multiplicity of plasmid transformations in *E. coli* has previously been shown to correlate with the DNA concentration used during a transformation (6, 8). Thus, to validate our dual-barcode approach, we created a small pUC19 plasmid library, containing the dual-barcoded region, which we then transformed into the *E. coli* cloning strain TOP10 (Fig. 5A-B). We added 1, 10 or 100 ng of our plasmid DNA library to each electroporation reaction. Post-electroporation, we colony PCR amplified the barcoded region from all plasmids found within 31 individual single-cell-derived bacterial colonies per experiment. PCR amplified DNA, containing the barcoded region of each pUC19 plasmids, were Sanger sequenced and manually curated for the presence of multiple SNPs. From this analysis we observed that when our TOP10 cells were electroporated with 1 ng of plasmid DNA, then 93% of all colonies contained one unique plasmid sequence (Fig. 5A). While only 58% and 42% of cells were mono-transformed when 10 ng or 100 ng of DNA were used, respectively (Fig. 5A). However, while 1 ng yielded the highest quality library, it also resulted in >1-log fewer transformed bacterial cells per reaction compared to when 10 ng or 100 ng of plasmid DNA was electroporated (Fig. 5B). These results support previous findings, which demonstrated that increasing extracellular DNA concentrations during electroporation of *E. coli* also increases transformation yields, yet at the expense of decreasing the fraction of mono-transformed cells (8).

**Figure 5.**
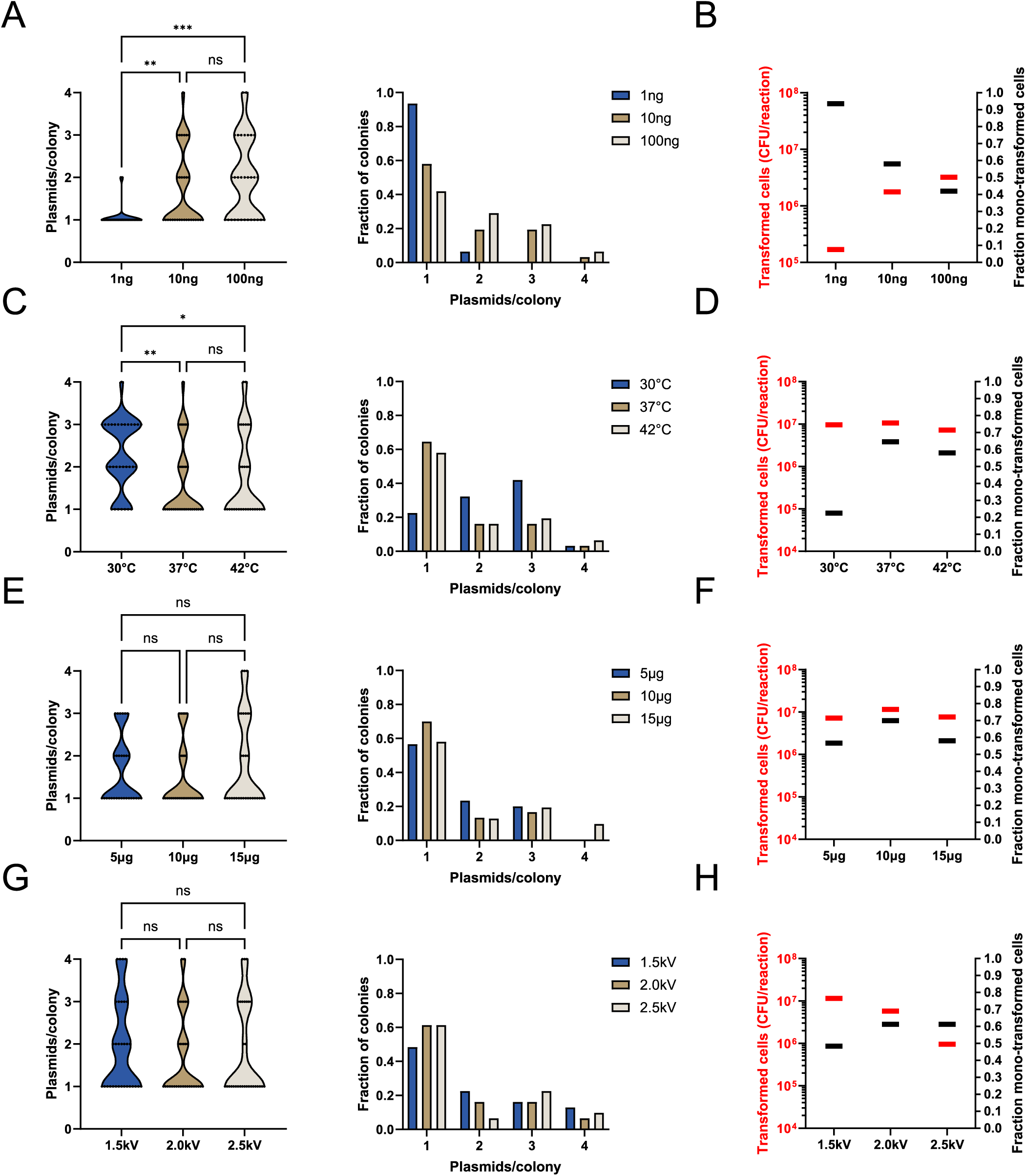
A mild heat-treatment before electroporation increases the fraction of mono-transformed yeast cells, without diminishing transformation yields. A) Plasmid enumeration of single-cell derived *E. coli* TOP10 bacterial colonies, transformed with different concentrations of a pUC19 dual-barcoded plasmid library (n=31). B) Correlation between transformation yields and the fraction of mono-transformed *E. coli* cells during an electroporation, as a result of the plasmid DNA concentration used. Fraction of mono-transformed cells was calculated from data in panel A. C-H) Plasmid enumeration of single-cell derived *S. cerevisiae* strain CEN.PK110-10C transformed with a pRS413 dual-barcoded plasmid library, using the HEEL method. Changing only the C) heat-shock temperature (n=31), E) plasmid DNA concentration (n=30-31), or G) electroporation voltage (n=31). Correlation between transformation yields and the fraction of mono-transformed yeast cells, as a result of the D) heat-shock temperature, F) plasmid DNA concentration, or H) electroporation voltage used. Fraction of mono-transformed yeast cells was calculated from data in panel C, E and G, respectively. Statistical significance was calculated by Two-way ANOVA with ns: P>0.05, *: P≤0.05, **: P≤0.005 and ***: P≤0.0005.

### A mild heat-treatment before electroporation increases the fraction of mono-transformed yeast cells without diminishing transformation yields

Armed with the knowledge that our dual-barcode approach allowed for a fast and efficient quantification of the multiplicity of DNA transformations, we recombined our pUC19 plasmid library with a pUC19-based yeast shuttle vector (pRS413) *in vivo* and then proceeded to transform our CEN.PK110-10C model strain with our new pRS413 yeast plasmid library. As we already observed that treating yeast cells with a mild 37°C heat-shock prior to electroporation resulted in no reduction in the transformation yield, even though fewer cells survived the treatment, we were curious to investigate if the increased fraction of surviving transformed cells would also translate to an effect on the multiplicity of transformation (Fig. 2F). Thus, we subjected our CEN.PK110-10C yeast cells to either a 42°C or 37°C heat-treatment, or a 30°C incubation, before electroporation. We then isolated and colony PCR amplified the plasmid-encoded barcodes from 31 individual single-cell derived yeast colonies per condition (Fig. 4A). All 93 yeast colony PCR reactions succeeded in generating an easy to sequence PCR product. From the Sanger sequencing chromatograms, we were able to conclude that approximately 15-25% of our plasmid library consisted of vectors that lacked >3 nt of the barcoded region, and approximately 10-15% of the library contained small 1-3 nt deletions within the barcoded sequence. Using a combination of both the SNP and the high-diversity region within the dual-barcode we were still able to quantify the multiplicity of plasmid transformations for all 93 colonies (Fig. 4B). Similar to previous findings by Scanlon *et al.*, we could also observe that when we used our improved yeast electroporation protocol, and when transformation yields were high (>10^7^ transformed cells per reaction), only 20% of all cells were mono-transformed (Fig. 5C) (7). However, we could also observe that a mild 37°C heat-shock prior to electroporation drastically increased the fraction of mono-transformed cells, with almost 70% of all transformed yeast cells being mono-transformed (Fig. 5C). Heat-shocking cells at a higher temperature (42°C) did not have an additional beneficial effect. Furthermore, as this enrichment of mono-transformed cells through heat-shocking did not reduce the transformation yields, we concluded that heat-shocking yeast cells before electroporation is a fast, efficient, and exceptionally convenient way of increasing the quality of a transformed yeast DNA library, without sacrificing diversity (Fig. 5D). Next, we were curious to investigate if changing the DNA concentration or electroporation voltage could increase this effect even further. Remarkably, neither the fraction of mono-transformed cells or the transformation yield changed significantly when we heat-shocked cells at 37°C, but either decreased or increased the amount of plasmid DNA to 5 µg or 15 µg, respectively (Fig. 5E-F). Likewise, changing the electroporation voltage to either 1.5 kV or 2.5 kV did not significantly change the fraction of mono-transformed cells (Fig. 5G). Although, the transformation yield did decrease by more than 1-order of magnitude when a voltage of 2.5 kV was used, compared to either 1.5 kV or 2.0 kV (Fig. 5H). Leaving us to conclude that our mild heat-shock and electroporation method could not easily be improved further by changing the physical and chemical parameters investigated in this study. Considering these findings, we named our new yeast transformation methodology **He**at-Shock **El**ectroporation (HEEL) (Fig. 3A).

### MiSeq sequencing validates the dual-barcode- and Sanger-based plasmid enumeration approach

To validate our approach of using a dual-barcode and simple Sanger sequencing to quantify the multiplicity of plasmid transformation(s) in yeast, we designed our automated yeast colony PCR workflow to also apply Illumina MiSeq P5 and P7 flow-cell binding adapter sequences, along with unique, experiment/condition specific, Illumina i7 barcodes (Fig. 4A). This allowed the same DNA fragments sequenced by Sanger sequencing to also be sequenced by Illumina MiSeq NGS. After the number of unique plasmids was enumerated by conventional Sanger sequencing, the DNA from all 279 individual yeast colony PCR reactions (31 colonies x 9 conditions), excluding 2 failed PCR reactions, was pooled, purified, and sequenced using the Illumina MiSeq platform. From the Sanger sequencing data, we knew that we had a low-diversity library of <1,000 total unique sequences in our pooled MiSeq library. Thus, we hypothesized that with our low-diversity library, and the high sequencing coverage we would achieve with an NGS approach, we would regrettably detect even rare mutagenic events and/or sequencing error(s) at a high level, which would lead us to erroneously identify unique barcoded sequences. Some of the possible sources of errors during our NGS analysis would include, but not be limited to, cross-well contamination(s) during the colony PCR sample preparation (a common problem for unspecific metagenomic sequencing (14)), mutations introduced by the Taq DNA polymerase during the PCR amplification of the barcoded region (8 x 10^-6^ mutations/bp/DNA duplication (15)), MiSeq sequencing errors (0.2-1.2% chance of incorrect base calling, dependent on the sequence context (16, 17)), as well as index hopping during the NGS demultiplexing (0.2-6% (18)). In the case of index hopping and cross-well contaminations during sample preparations, this would lead to unique barcodes being assigned to the wrong library (experimental conditions), and thus falsely inflate the actual plasmid count for each experimental condition. PCR-mutations and MiSeq-dependent sequencing errors would both lead to the creation of additional unique barcoded sequences for each experimental condition, which would also falsely inflate the actual plasmid count per condition. To overcome these issues, we processed, filtered, and curated our NGS data in three steps (Fig. 6A). **1)** First, we removed all sequencing reads that had an average base-calling quality of <Q35. This removed 15-20% of all reads/experimental condition. **2)** Then we chose to apply a rational cut-off of 300-fold coverage per unique barcode to best exclude erroneous sequences and remove them from our downstream NGS analysis. The 300-fold coverage was selected as we observed that a large proportion of unique barcodes had a very low abundance of only 10- to 100-fold coverage (Fig. S2). We concluded that these low-abundant barcodes most likely constituted our PCR, and/or NGS-specific background. **3)** Finally, we performed a sequence similarity analysis where we curated and designated identical, or near-identical sequences (only 1 nt difference) found in multiple libraries as “uncertain”. I.e., if two identical sequences occurred in 2 or more libraries (experimental conditions), then one of these sequences would most probably be the result of an index hopping error, or from a cross-well contamination event during the sample preparation. Likewise, if two sequences were near-identical, i.e., only differed by 1 nt, then we reasoned that one of these sequences most likely was the results of a PCR or MiSeq sequencing error. The rationale behind this hypothesis was as follows; each dual-barcode sequence can create a maximum of 3.46 x 10^7^ unique sequences (1,048,576 (10N) x 33 possible SNPs), or a minimum of 8.65 x 10^6^ unique sequences (262,144 (9N) x 33 possible SNPs), if 1 nt is allowed to be identical (Fig. S1A). Since we only expected <1,000 random sequences in our NGS data, the likelihood that two unique sequences, or near-identical sequences, simultaneously occurring in multiple libraries/experiments is highly unlikely. However, as we could not determine if these ambiguous sequences should be excluded, because they arose from index hopping or cross-well contamination, or if two similar sequences should be counted as one, if they arose from PCR or sequencing errors, we instead choose to use these uncertain sequences to calculate the range of the most probable number of plasmids per library (Table 1). After we curated our NGS data, we could clearly verify our previous findings, as the MiSeq NGS and Sanger based measurements closely correlated with each other (Table 1). For 7 of the 9 experimental conditions, the Sanger-based plasmid enumeration was within the range of the most probable number of plasmids calculated using the NGS data (Table 1). For library 3 (42°C heat-shocked cells), the minimum number of NGS enumerated unique plasmid sequences was approx. 15% higher than the number of unique plasmids enumerated by Sanger sequencing. While for library 9 (2.5 kV electroporation) this discrepancy was only approx. 7% (Table 1). Lastly, we analyzed if our random dual-barcode library contained any sequence bias that might affect our NGS analysis (Fig. 6B). From this analysis we could conclude that all 4 nucleotides could be found at approximately equal abundance at each of the 10 random positions, and in the 11 nt SNP-based barcode we could observe that all 3 alternative nucleotides existed at every position of the barcode, although at a low abundance (Fig. 6B). As we expected 1 SNP/barcode, and there were 33 possible SNP combinations, this low abundance clearly demonstrated that our barcoding workflow had produced the intended diversity, nucleotide distribution and sequence composition. Collectively, from these findings we concluded that our NGS data principally correlated with our Sanger sequencing results and that our dual-barcoding approach allowed for an accurate enumeration of the number of plasmids/yeast cell, in the ≤4 plasmids/yeast cell range (Table 1). We could also conclude that regardless of the plasmid enumeration deviation for library 3 and 9, our Sanger sequencing and NGS data independently led to the same experimental conclusions, i.e., that a mild heat shock (37°C) reduced the multiplicity of plasmid transformations in yeast without diminishing the transformation yield.

**Figure 6.**
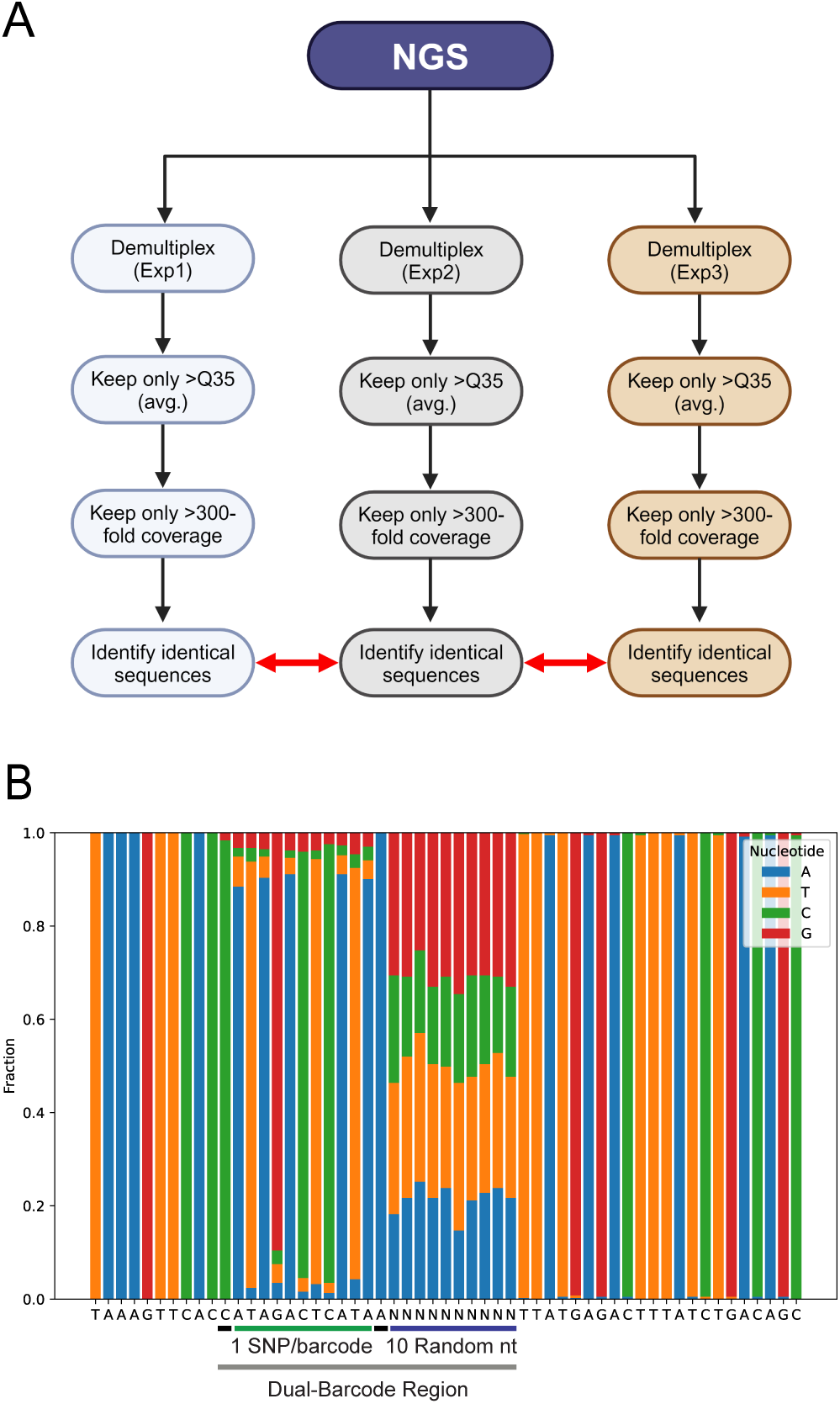
Overview of the NGS workflow. A) Schematic illustration of the NGS workflow used to enumerate the most probable number of unique dual-barcoded sequences in each library (experimental condition). Illustration created with BioRender.com. B) Sequence bias and nucleotide frequency analysis of the dual-barcode region. The analysis includes all NGS-identified dual-barcoded sequences with the entire barcoded region intact.

**Table 1.** MiSeq sequencing validates the dual-barcode and Sanger-based plasmid enumeration approach. Comparison of the Sanger and NGS derived plasmid enumerations, including the occurrence of identical and near-identical (1 nt difference) sequences within and between different libraries (experimental condition). Uncertain sequences were subtracted from the NGS enumerated sequences to obtain a range of the most probable nr of plasmids per library.

| Library | Condition | Sanger Sequencing | NGS |  |  |  |  |
| --- | --- | --- | --- | --- | --- | --- | --- |
|  |  | Total nr of plasmids | Total nr of plasmids | Nr of identical sequences | Nr of near-identical sequences | Uncertain sequences | Most probable nr of plasmids |
| Exp1 | 30°C (No heat-shock) | 70 | 94 | 22 | 3 | 25 | 69-94 |
| Exp2 | 37°C (Mild heat-shock) | 49 | 58 | 15 | 0 | 15 | 43-58 |
| Exp3 | 42°C (Heat-shock) | 54 | 79 | 16 | 1 | 17 | 62-79 |
| Exp4 | 5µg plasmid DNA | 49* | 56* | 18 | 0 | 18 | 38-56 |
| Exp5 | 10µg plasmid DNA | 44* | 49* | 12 | 0 | 12 | 37-49 |
| Exp6 | 15µg plasmid DNA | 56 | 62 | 14 | 1 | 15 | 47-62 |
| Exp7 | 1.5kV electroporation | 60 | 70 | 17 | 0 | 17 | 53-70 |
| Exp8 | 2.0kV electroporation | 52 | 58 | 14 | 1 | 15 | 43-58 |
| Exp9 | 2.5kV electroporation | 56 | 77 | 16 | 1 | 17 | 60-77 |
| Total number of plasmids |  | 490 | 603 | 144 | 7 | 151 | 452-603 |

\* = Result was calculated from 30 individual yeast colonies, rather than 31.

## Discussion

Multiplicity of plasmid transformations in bacteria and yeast is a well-known problem, but also a scarcely investigated area of research. To the best of our knowledge, no study has designed any mitigation strategies to address this issue, beyond merely lowering the concentration of DNA used during a transformation. Though, this is a highly counterproductive strategy, as decreasing the concentration of DNA during a transformation also decreases the number of transformed cells (transformation yield). I.e., there is an inverse correlation between the fraction of mono-transformed cells and the transformation yield. Here we have demonstrated an easy and straight-forward yeast transformation method that significantly increases the fraction of mono-transformed yeast cells within a population, without sacrificing the transformation yield. Our strategy relies on exposing yeast cells to a mild, 37°C heat-shock, before electroporating the cells with both plasmid and single-stranded salmon sperm DNA. Using this strategy (HEEL) we can increase the fraction of mono-transformed cells from 20% to 70%, while still maintaining a high transformation yield of approx. 2 x 10^7^ transformed yeast cells per reaction. Thus, we can significantly increase the number of easy-to-screen high-quality yeast transformants with clear phenotype-to-genotype relationships, while still retaining a high-diversity library. Current methods are restricted to generating either high-diversity libraries with low-resolution (multiple genotypes per cell), or high-resolution libraries, but with a low-diversity (few transformed cells). We expect that the HEEL methodology will simplify DNA, RNA and/or protein library screenings in yeast to such an extent that high-diversity and high-resolution libraries can be generated, concurrently, with ease.

To the best of our knowledge, no study has reported that heat-shocking yeast cells before electroporation can elicit a beneficial effect on DNA transformations. Presumably this is because most studies have only investigated the effect that a traditional heat-shock temperature (42°C) has on yeast cells. And, as we show here, such high temperatures only have a detrimental effect on yeast plasmid uptake during an electroporation. Also, since a mild heat-shock temperature of 37°C does not appear to affect transformation yields in any significant way it is easy to understand why this effect might have been overlooked by others. Although, as we show here, a mild heat-shock before electroporation does increase the fraction of transformed yeast cells, indicating that the efficiency of DNA uptake is increased, simultaneously as more cells die by the heat-shock. Looking towards previously presented yeast DNA transformation models, it is possible that a novel synergistic effect might be happening. I.e., that a membrane permeability effect, mediated by the electroporation, is occurring simultaneously with an endocytosis effect, mediated by Lithium acetate, heat and ssDNA. However, this is of course merely speculation, and we have intentionally chosen not to attempt to decipher the exact molecular mechanism behind this effect in this work. Moreover, our results clearly support previous findings that Lithium acetate, DTT, ssDNA and elevated temperatures, are all factors of paramount importance for a successful yeast DNA transformation (2, 3, 5, 12). Although, we observe that these four parameters can, under some conditions, give rise to novel and unique transformation effects when applied in a nonconventional way, i.e., by using heat-shock and electroporation simultaneously.

In regard to our inability to reproduce the findings of Benatuil *et al.*, we observed that less than 0.2% of our EBY100 yeast cells survive the electroporation protocol described by Benatuil *et al.* (Fig. 1) (5). While Benatuil *et al.* reported to be able to transform >84% of all yeast cells in solution. Thus, there is a great discrepancy between our results presented here and the transformation yields reported by Benatuil *et al.* (5). This discrepancy has also been described by multiple other studies, which have all reported transformation yields of 10^6^-10^7^ transformants/reaction (19–21). This is approximately 10- to 100-fold lower than the 5.42 x 10^8^ transformants/reaction reported by Benatuil *et al.* (5). Although, it should be noted that the 10^6^-10^7^ transformants/reaction that were reported by others far exceeds the 10^3^ transformants/reaction we could achieve in our lab (Fig. 1) (5). A likely explanation for this could be that unlike Benatuil *et al.*, and other studies, we have not transformed linear DNA molecules, rather we have used super-coiled circular DNA. Since linear DNA with homology-arms has previously been shown to transform yeast better than both linear DNA, without homology-arms, and circular DNA, it is likely that DNA topology issues is the reason for our reduced DNA transformation efficiency (22). However, this hypothesis still does not explain why only 0.1-0.2% of the EBY100 yeast cells survive the Benatuil *et al.* electroporation method in our laboratory (Fig. 1) (5). Nevertheless, our findings presented here clearly demonstrate that by modifying the method described by Benatuil *et al.*, by lowering the electroporation voltage from 2.5 kV to 2.0 kV and by adding 100 µg of single-stranded salmon sperm DNA to each transformation reaction, we can overcome this possible DNA topology issue and increase the transformation yield by almost 100-fold. No other modifications were necessary to successfully transform yeast cells with a circular DNA molecule and achieve a high-throughput transformation yield that is comparable to other studies that have transformed yeast with linear DNA molecules (19–21). Consequently, with these innovations we have made it possible to easily transform yeast cells with high-diversity libraries that first have been cloned and transformed into *E. coli.* This overcomes a major bottle-neck for most yeast library transformations, as having enough DNA to transform a high-diversity library in yeast is not always guaranteed. The creation of high-diversity libraries in yeast requires >5 µg of DNA, while only <50 ng is required for an efficient DNA transformation of *E. coli* (1, 4, 6). By using the HEEL method described here, it is possible to first transform a high-diversity library into *E. coli*, allow the bacteria to amplify the plasmid DNA library as much as needed, then extract the plasmid DNA and immediately transform this super-coiled circular plasmid DNA library into yeast. Here, we completely omit the need to convert the plasmid to a linear molecule with homology-arms to achieve high-efficiency transformations.

The development of NGS techniques has revolutionized the quantification and sequencing of high-diversity mutant libraries. However, the inherent weakness of any high-throughput NGS technique lies in its inability to link a single-cell phenotype to a specific genotype. A diverse library of mutants is often first screened for novel-function(s), then each cell that exhibits a desired phenotype is selected/sorted and then pooled/binned together with all other cells with a similar phenotype. These mutants are then sequenced as a group to identify each individual genotype found in this sorted library. While this strategy generates a large pool of sequence data, it does not allow for a precise phenotype-to-genotype determination, merely an approximate phenotype-to-genotype relationship, as the range of different cellular phenotypes in each sorted library can differ by as much as one order of magnitude. In the past, this has usually not been a source of concern, as most mutant screening endeavors were designed to merely identify all beneficial mutants, of which a few clones were then chosen for further screening and characterization. However, with the introduction of AI and machine learning algorithms in most scientific fields, this is no longer the case. Predictive algorithms require precise phenotype-to-genotype measurements to generate insightful models with high-accuracy. Approximate phenotype-to-genotype measurements are non-optimal as an AI training set. To create better AI generated models, we need to strive to generate better data with better resolution. As shown here, the HEEL methodology can minimize the problem of phenotypic noise, i.e., having multiple genotypes present in each cell during a mutant yeast screening. Our automated genotyping workflow allows for hundreds of yeast cells to be efficiently genotyped, which significantly simplifies the identification of beneficial mutants when using yeast as a screening platform (9). In addition, while the number of mutants screened and genotyped would be limited to just a few hundred, predictive algorithms should nevertheless be able to generate more insightful models from this smaller, but more high-resolution data. This new approach should avoid the issues of binning sorted cells based on phenotype and then using the average phenotype of the group to perform a linear regression, which can lead to incorrect conclusions and mathematical models, as intrinsic noise in the data is overlooked (23).

Furthermore, while our dual-barcode approach can only be used to enumerate a maximum of 4-6 unique sequences per cell, the SNP-based barcode, by itself, can also be used to quantify specific genotypes in a mixed population. Our 11 nt SNP-based barcode can create 33 individual combinations (11 nt x 3 alternative nucleotides per position). Each of these 33 genotypes can, for example, be assigned to a specific species, strain or protein variant and thus be used as a unique molecular identifier during a Sanger sequencing reaction. Since each individual SNP can be identified precisely during a Sanger sequencing, this approach can be used to genotype a mixed population of <6 different strains/constructs per cell using only one single sequencing reaction. Thus, this approach is expected to be highly valuable when screening combinatorial libraries with a low-diversity. Such libraries would include, but not be limited to, the identification of the optimal combination of promotor(s), terminator(s) and/or vector backbone(s) to achieve optimal recombinant protein expression in a new host, although many other custom applications for such a SNP-based genotyping approach could be designed.

## Material and Methods

### Strains, DNA constructs and growth conditions

The bacterial and yeast strains, plasmid library and oligos used in this study are listed in Tables S1– S3. A detailed description of the dual-barcode plasmid library construction is available in the supplementary material and methods section of the appendix, found in the online version of this article. Bacterial strains were routinely grown at 37°C and with shaking at 200 r.p.m. in lysogeny broth (LB): 10 g/l tryptone, 5 g/l yeast extract and 10 g/l NaCl. Media was supplemented with ampicillin, at 100 mg/l, or Chloramphenicol, at 25 mg/l when appropriate. Yeast strains were routinely grown at 30°C and with shaking at 200 r.p.m. in yeast peptone dextrose broth (YPD): 20 g/l peptone, 10 g/l yeast extract and 20 g/l Glucose, or in synthetic drop-out medium, minus leucine (SC-Leu), (Sigma-Aldrich, #Y1376), or minus histidine (SC-His), (Sigma-Aldrich, #Y1751).

### *E. coli* DNA electroporation protocol

100 mL of a *E. coli* culture were grown to OD600 = 0.4 then centrifuged at 3000xg for 10min. The resulting pellet was pooled and re-suspended in 50 mL ultra-pure Milli-Q water. This procedure was repeated 2 more times and the triple washed cell pellet was then suspended in a total volume of 400 µL Milli-Q water to achieve a final cell concentration of approximately OD600 = 100. 50 µL of this concentrated cell solution and 1, 10 or 100 ng of plasmid DNA was transferred to a 1 mm MicroPulser electroporation cuvette (Bio-Rad) and cells were then electroporated at 1.8 kV, 25 µFD and 200 Ohm, using a Gene Pulser Xcell Electroporation System (Bio-Rad). 500 µL of LB was added and cells were left to recover for 90min at 37°C before being inoculated into LB-both or onto LB-agar with appropriate antibiotic selection.

### Heat-Shock Electroporation (HEEL) yeast DNA transformation protocol

A detailed description of the HEEL yeast DNA transformation method can be found in the supplementary material and methods section of the appendix, found in the online version of this article. For the preliminary HEEL method optimization (Fig. 2A-F), the transformation protocol was carried out exactly as the optimized HEEL protocol (Fig. 3A) except that yeast cells were conditioned in a solution of 100 mM Lithium acetate and 20 mM Dithiothreitol (DTT), at either 30°C or 42°C for 1h. Cells were also electroporated with only 5 µg of plasmids DNA and 100 µg of salmon sperm ssDNA, unless otherwise stated.

### Automated yeast genotyping workflow

A detailed description of the automated yeast genotyping workflow can be found in the supplementary material and methods section of the appendix, found in the online version of this article.

### MiSeq NGS sequencing

Yeast single-cell derived colony PCR amplified amplicons were pooled according to experimental conditions, purified using a silica DNA-binding column, and then sequenced with a MiSeq Reagent Micro Kit v2 (300 Cycles) on an Illumina MiSeq instrument, according to manufacturer’s instructions. The pooled libraries were sequenced at a concentration of 10 pM, with 12.5 pM PhiX Control v3 (Illumina) added. A custom read primer (oligo 358) and a custom index primer (oligo 359) were used at a final concentration of 0.5 µM during the MiSeq sequencing to overcome potential issues with primer-dimer fragment sequencing (Table S3).

### NGS analysis

A detailed description of the Illumina NGS sequencing analysis can be found in the supplementary material and methods section of the appendix, found in the online version of this article.

## Acknowledgements

This work was supported by the Wenner-Gren Foundations, to MW, and The Novo Nordisk Foundation, grant number NNF20SA0035588, to EEHS, NNF210C0069089 and NNF20CC0035580, to SG, and NNF20CC0035580, to MDJ and EDJ. The authors would like to thank the DNA Foundry at The Novo Nordisk Foundation Center for Biosustainability for supporting the HEEL NGS sequencing project.

## Author contributions

**Marcus Wäneskog**: Conceptualization; Designed research; Performed research; Analyzed data; Funding acquisition; Wrote the paper – original draft, review(s) and editing. **Emma Elise Hoch-Schneider:** Performed research; Analyzed data. **Shilpa Garg:** Analyzed data. **Christian Kronborg Cantalapiedra:** Analyzed data. **Elena Schaefer:** Performed research. **Michael Krogh Jensen:** Project administration; Resources; Funding acquisition; **Emil D. Jensen**: Project administration; Resources; Funding acquisition; Wrote the paper – review(s) and editing.

