## Supplementary appendix for "Heat pre-treatment reduces multiplicity of plasmid transformations in yeast during electroporation, without diminishing the transformation efficiency"

**This supplementary file includes:**

Supplementary methods

Supplementary tables S1-S3

Supplementary figures S1-S2

**Yeast Heat-Shock Electroporation (HEEL) DNA transformation workflow**

60 mL YPD broth (1% (w/v) yeast extract, 2% (w/v) meat-derived peptone and 1% (w/v) glucose) was inoculated with a fresh re-streaked yeast colony and incubated at 30°C with vigorous aeration for 24h, or until the yeast culture reached stationary phase.

40 mL of the stationary phase yeast culture was then added to 400 mL of fresh YPD broth and incubated at 30°C with vigorous aeration for 3-4h, until reaching OD600 = 1.

400 mL of the OD600 = 1 yeast culture was collected and centrifuged at 3000xg for 5min. The YPD supernatant was discarded, and the resulting yeast cell pellet was collected and pooled in a total volume of 50 mL pre-chilled (4°C) ultrapure Milli-Q Water. This washing step was repeated one additional time.

The resulting yeast pellet was then resuspended in 50 mL of a 1 M sorbitol and 1 mM CaCl2 solution.

The osmotically shocked yeast cells were then centrifuged again at 3000xg for 5min, and the sorbitol solution was discarded, and the yeast cells were resuspended in 50 mL of a 100 mM Lithium Acetate (LiAc) and 10 mM Dithiothreitol (DTT) solution. The LiAc and DTT solution was then incubated at 37°C with vigorous aeration for 1h.

The mildly heat-shocked cells were then pelleted at 3000xg for 5min and resuspended in 50 mL of a 1 M sorbitol and 1 mM CaCl2 solution. After an additional centrifugation at 3000xg for 5min, the pelleted yeast cell solution was re-suspended in a 1 M sorbitol and 1 mM CaCl2 solution, in a total volume of 4 mL.

400 µL of the yeast solution was used per transformation reaction by mixing the cell suspension with 10µg of circular plasmid DNA and 100µg of boiled single-stranded salmon sperm DNA (ssDNA). The plasmid DNA and ssDNA mixture was then transferred to an MicroPulser electroporation cuvette (Bio-Rad) with a 2 mm gap. The solution was immediately electroporated at 2.0 kV, 200 Ohm and 25µFD, using a Gene Pulser Xcell Electroporation System (Bio-Rad). This resulted in a time constant of approximately 3.5-4.2 ms. Directly after the electroporation (within 5sec), 2 mL of fresh YPD was added to the electroporated cells and the solution was incubated at 30°C with vigorous aeration for 1h to allow cells to recover.

Starting/input cells, surviving cells and transformed cells were all enumerated by first serial diluting the starting cell solution and the recovered cell suspension. Each dilution was inoculated on a YPD (surviving cells) and selective media (transformed cells) agar plate. Yeast colonies were enumerated after 48h of growth at 30°C.

**Automated yeast genotyping workflow**

100 µL of a yeast cell wall digestion solution, containing 10 mM TRIS-HCl pH 8, 1 mM Ethylenediaminetetraacetic acid (EDTA), 10 mM Dithiothreitol (DTT) and 1000 U/mL Lyticase from Arthrobacter luteus (Sigma- Aldrich), were dispensed equally into a 96-well plate (Thermo Fisher Scientific) using an OT-2 liquid handling robot (Opentrons).

Individual single-cell derived yeast colonies were randomly picked using a Singer PIXL Precision Microbial Colony Picker (Singer Instruments) and inoculated into each of the pre-aliquoted 96 wells, using the following settings:

Pinning Dry (Speed: 19 mm/s, Pickup Line Length: 10 mm)
Mixing Dry (Speed: 20 mm/s, Radius: 0.4 mm, Vertical Travel: 2.5 mm, Cycles: 1)
Pinning Wet (Speed: 60 mm/s, Well Base Clearance: 20% of well, Nozzle Clearance: 2 mm)
Mixing Wet (Speed: 60 mm/s, Radius: 50% of available, Revolutions: 3)

The Lyticase and yeast colony mixture were incubated at 37°C for 1 hour. Then 90 µL of the mixture was removed and discarded by an OT-2 Robot (Opentrons). The remaining 10 µL was mixed with 14 µL of a sterile MilliQ $H_2O$ solution containing a universal forward primer (oligo; 348).

9 separate RedTaq DNA polymerase master mixes (Sigma- Aldrich), one per experiment, was prepared and mixed with experiment specific reverse primers (oligos; 349-357). 26 µL of each PCR master mix was added to each of the yeast cell solutions, using an OT-2 Robot (Opentrons). The 96-well plate was then incubated in a T100 Thermocycler (Bio-Rad), using the following settings:

1) 95°C for 10 min

   2) 95°C for 25sec
   3) 61°C for 30sec
   4) 72°C for 30sec.

   Step 2-4 was repeated for 35 cycles.

5) 72°C for 5min

Following the PCR amplification, 15 µL from 31 individual PCR reactions per experiment was transferred to a 96-well sequencing plate, leaving 3 empty wells as negative controls, and Sanger sequenced (Eurofins Genomics) using oligo; 288. The remaining 35 µL of each PCR reaction was purified and pooled for Illumina MiSeq NGS and analysis.

**MiSeq NGS analysis workflow**

The NGS analysis was performed in Python using the following libraries: Biopython, matplotlib, Collections, Pandas. The following steps outline the data processing:

*Sequence Filtering by Phred score:*
Sequences from FASTQ files were parsed using Biopython, and each sequence's average Phred quality score was computed. Sequences with an average quality score below a defined threshold (Q35 in this study) were discarded. This filtering step ensured that only high-quality sequences were retained for further analysis.

*Alignment to Reference Sequence:*
The filtered sequences were aligned to a reference sequence (TGCGCCAGCTTTCATCCCCGATATGCACCACCGGGTAAAGTTCACCATAGACTCATAANNN NNNNNNNTTATGAGACTTTATCTGACAGCAGACGTGCACTGGCCAGGGGGATCACCATCC GTCGCCCGGG) using a global pairwise alignment with a gap penalty of -1. This alignment treated 'N' as either A, C, T, or G and sequences were then trimmed to a predefined region for uniform length (position 36 to 91 of the reference sequence).

*Sequence Counting:*
After the reference sequence alignment, all unique sequences were identified and counted in each of

the 9 experimental libraries. Sequences with a coverage of more than 300-fold were retained for further analysis.

*Visualization and Reporting:*
The processed data, including the counts of unique sequences and nucleotide frequencies, was reported and visualized with matplotlib. For plotting of nucleotide frequencies, all sequences with indel, compared to the reference sequence, were excluded to visualize the bias within the dual barcode region.

**Construction of the pRS413-CEN/ARS-ccdB-cat plasmid**

The 2-micron origin of replication of the previously described pESC-HIS-ccdB-USER (pCfB55) plasmid was changed by yeast *in vivo* recombination by first cutting the 2μ origin of replication with FD XbaI (Thermo Fisher Scientific), thus linearizing the plasmid (1). The previously described pRS413 plasmid, which has a CEN/ARS origin of replication, but otherwise share an identical backbone to the pCfB55 plasmid, was linearized by cutting-out a fragment containing the CEN/ARS origin of replication flanked by partial AmpR and HIS3 CDS sequences, using FD HindIII (Thermo Fisher Scientific) and BsrDI (New England Biolabs) (2). The pRS413 fragment containing the CEN/ARS origin of replication and the linearized pCfB55 plasmid was co-transformed into CEN.PK110-10C (MW2) using the lithium acetate ssDNA heat-shock method to allow for homologues recombination of the two fragments (3). Yeast colonies that were histidine prototrophs were PCR screened using oligos; 73 and 17031, to identify positive recombination events. The recombined yeast plasmid (pMW1) was extracted using the Zymoprep Yeast Plasmid Miniprep Kit (Zymo Research) and transformed into a CcdB resistant *E. coli* strain (MW69) (Thermo Fisher Scientific). Following a plasmid extraction from *E. coli*, the entire plasmid was verified by whole plasmid sequencing using Oxford Nanopore Technologies (ONT).

**Construction of the Dual-Barcoded pUC-ccdB-cat plasmid**

The pUC19 backbone along with the chloramphenicol resistance marker and the CcdB counterselection marker was PCR amplified with oligos; 141 and 142, from the pRS413-CEN/ARS-ccdB-cat plasmid (pMW1). The resulting linear DNA was self-circularized, using T4 DNA ligase (Thermo Fisher Scientific), and then transformed into a CcdB resistant *E. coli* strain (MW69) (Thermo Fisher Scientific). Plasmids from several chloramphenicol resistant bacterial colonies were extracted and verified by sequencing, using oligo; 237. This new pUC19-CatR-CcdB vector (pMW17) was then used as template for a PCR reaction using a mix of 12 individual oligos (272-283). Introducing the dual-barcoded sequences into the middle of the ccdB CDS. The resulting linear DNA was self-circularized at a final concentration of 1 ng/μL, using T4 DNA ligase (Thermo Fisher Scientific). This circular molecule was transformed via electroporation into the CcdB sensitive *E. coli* strain TOP10 (MW150) (Thermo Fisher Scientific), selecting for plasmids with a ccdB gene that was disrupted by an insertion of the dual-barcoded sequence. Plasmid selection and propagation post-transformation was done entirely in liquid culture to ensure maximum diversity. The entire plasmid library (pMW35) was extracted and sequenced with oligo; 288, to verify diversity and the presence of the dual-barcode.

#### Construction of the Dual-Barcoded pRS413-CEN/ARS-ccdB-cat plasmid

The pRS413-CEN/ARS-ccdB-cat plasmid (pMW1) was linearized by cutting in the middle of the ccdB CDS with FD SmaI (Thermo Fisher Scientific). This linear pRS413-CEN/ARS-ccdB-cat fragment was then co-transformed into CEN.PK110-10C (MW2) along with the Dual-Barcoded pUC-ccdB-cat plasmid library (pMW35), using the HEEL yeast DNA transformation methodology. This allowed the Dual-Barcoded pUC-ccdB-cat library to be used as a gap-repair template for the linear pRS413-CEN/ARS-ccdB-cat fragment, ensuring a recombination of the dual-barcoded region from the pUC-ccdB-cat plasmids to the pRS413-CEN/ARS-ccdB-cat plasmid. The resulting yeast plasmid library (MW165) was then extracted using the Zymoprep Yeast Plasmid Miniprep Kit (Zymo Research) and transformed into the CcdB sensitive *E. coli* strain TOP10 (MW150) (Thermo Fisher Scientific). Selecting for pRS413-CEN/ARS-ccdB-cat plasmids with a disrupted ccdB gene. The resulting counter-selected Dual-Barcoded yeast plasmid library was then amplified in TOP10 *E. coli* cells (pMW39) before being extracted and sequenced with oligo; 288, to verify sequence diversity and the presence of the dual-barcode region.

**Table S1. Bacterial strains used in this study.**

| Strain number | Genotype | Origin |
|---|---|---|
| MW69 | *E. coli* One Shot ccdB Survival 2 T1R *F- mcrA Δ(mrr-hsdRMS-mcrBC) φ80lacZΔM15 ΔlacX74 recA1 araΔ139 Δ(ara-leu)7697 galU galK rpsL(StrR) endA1 nupG fhuA::IS2* | Thermo Fisher Scientific |
| MW150 | *E. coli* TOP10 *F- mcrA Δ(mrr-hsdRMS-mcrBC) φ80lacZΔM15 ΔlacX74 recA1 araD139 Δ(ara-leu)7697 galU galK λ–rpsL(StrR) endA1 nupG* | Thermo Fisher Scientific |
| pMW1 | *E. coli* One Shot ccdB Survival 2 T1R *F- mcrA Δ(mrr-hsdRMS-mcrBC) φ80lacZΔM15 ΔlacX74 recA1 araΔ139 Δ(ara-leu)7697 galU galK rpsL(StrR) endA1 nupG fhuA::IS2* /pRS413(CEN6-ARS4 ccdB-cat) His+ AmpR CatR | This Study |
| pMW17 | *E. coli* One Shot ccdB Survival 2 T1R *F- mcrA Δ(mrr-hsdRMS-mcrBC) φ80lacZΔM15 ΔlacX74 recA1 araΔ139 Δ(ara-leu)7697 galU galK rpsL(StrR) endA1 nupG fhuA::IS2* /pUC19(ccdB-cat) CatR | This Study |
| pMW35 | *E. coli* TOP10 *F- mcrA Δ(mrr-hsdRMS-mcrBC) φ80lacZΔM15 ΔlacX74 recA1 araD139 Δ(ara-leu)7697 galU galK λ–rpsL(StrR) endA1 nupG* /pUC19(ccdB::(low/high-complexity-barcode)-cat) CatR | This Study |
| pMW39 | *E. coli* TOP10 *F- mcrA Δ(mrr-hsdRMS-mcrBC) φ80lacZΔM15 ΔlacX74 recA1 araD139 Δ(ara-leu)7697 galU galK λ–rpsL(StrR) endA1 nupG* /pRS413(CEN6-ARS4 ccdB::(low/high-complexity-barcode)-cat) His+ AmpR CatR | This Study |

**Table S2. Yeast strains used in this study.**

| Strain number | Genotype | Origin |
|---|---|---|
| MW2 | CEN.PK110-10C *MATα his3Δ1* | Lab Collection |
| MW23 | CEN.PK2-1C *MATa ura3-52 his3Δ1 leu2-3,112 trp1-289 MAL2-8c SUC2* | Lab Collection |
| MW71 | BY4741 *MATa his3Δ1 leu2Δ0 met15Δ0 ura3Δ0* | Lab Collection |
| MW165 | CEN.PK110-10C *MATα his3Δ1* /pRS413(CEN6-ARS4 ccdB::(low/high-complexity-barcode-library)-cat) His+ AmpR CatR | This Study |
| MW168 | EBY100 *MATa aga1::pGAL1-aga1-URA3 ura3-52 trp1 leu2-Δ200 his3-Δ200 pep4::HIS3 prbd1.6R can1 GAL* | Lab Collection |

**Table S3. Oligos used in this study.**

| Name | Sequence 5'-3' |
|---|---|
| 73 | GGAACAACACTCAACCCTATCTCG |
| 141 | 5'Phosphate-CCTTTTTGATAATCTCATGACC |
| 142 | GCAGCCTACTCGCTATTGTC |
| 237 | CTCTTTTGCTGACGAGAACAGG |
| 272 | 5'Phosphate-GTGAACTTTACCCGGTGGTGC |
| 273 | CATAHACTCATAANNNNNNNNNNNTTATGAGACTTTATCTGACAGCAGACG |
| 274 | CATAGBCTCATAANNNNNNNNNNNTTATGAGACTTTATCTGACAGCAGACG |
| 275 | CATAGADTCATAANNNNNNNNNNNTTATGAGACTTTATCTGACAGCAGACG |
| 276 | CATAGACVCATAANNNNNNNNNNNTTATGAGACTTTATCTGACAGCAGACG |
| 277 | CATAGACTDATAANNNNNNNNNNNTTATGAGACTTTATCTGACAGCAGACG |
| 278 | CATAGACTCBTAANNNNNNNNNNNTTATGAGACTTTATCTGACAGCAGACG |
| 279 | CBTAGACTCATAANNNNNNNNNNNTTATGAGACTTTATCTGACAGCAGACG |
| 280 | CAVAGACTCATAANNNNNNNNNNNTTATGAGACTTTATCTGACAGCAGACG |
| 281 | CATBGACTCATAANNNNNNNNNNNTTATGAGACTTTATCTGACAGCAGACG |
| 282 | CATAGACTCAVAANNNNNNNNNNNTTATGAGACTTTATCTGACAGCAGACG |
| 283 | CATAGACTCATBANNNNNNNNNNNTTATGAGACTTTATCTGACAGCAGACG |
| 288 | TCGCGGTGGCTGAGATCAGC |
| 348 | AATGATACGGCGACCACCGAGATCTACACTAGATCGCTCGTCGGCAGCGTCCATGC GCCAGCTTTCATCC |
| 349 | CAAGCAGAAGACGGCATACGAGATCGCTCAGTTCGTCTCGTGGGCTCGGCAGAGT GATATTATTGACACG |
| 350 | CAAGCAGAAGACGGCATACGAGATTATCTGACCTGTCTCGTGGGCTCGGCAGAGT GATATTATTGACACG |
| 351 | CAAGCAGAAGACGGCATACGAGATATATGAGACGGTCTCGTGGGCTCGGCAGAGT GATATTATTGACACG |
| 352 | CAAGCAGAAGACGGCATACGAGATCTTATGGAATGTCTCGTGGGCTCGGCAGAGT GATATTATTGACACG |
| 353 | CAAGCAGAAGACGGCATACGAGATTAATCTCGTCGTCTCGTGGGCTCGGCAGAGT GATATTATTGACACG |
| 354 | CAAGCAGAAGACGGCATACGAGATGCGCGATGTTGTCTCGTGGGCTCGGCAGAGT GATATTATTGACACG |
| 355 | CAAGCAGAAGACGGCATACGAGATAGAGCACTAGGTCTCGTGGGCTCGGCAGAGT GATATTATTGACACG |
| 356 | CAAGCAGAAGACGGCATACGAGATTGCCTTGATCGTCTCGTGGGCTCGGCAGAGT GATATTATTGACACG |
| 357 | CAAGCAGAAGACGGCATACGAGATCTACTCAGTCGTCTCGTGGGCTCGGCAGAGT GATATTATTGACACG |
| 358 | GATAACGGAGACCGGCACACTGGCCATATCG |
| 359 | CGTGTCAATAATATCACTCTGCCGAGCCCACGAGAC |
| 17031 | AGGGGTUCCGCGCACATTTCCCCGAAAAGT |

### Fig. S1

A

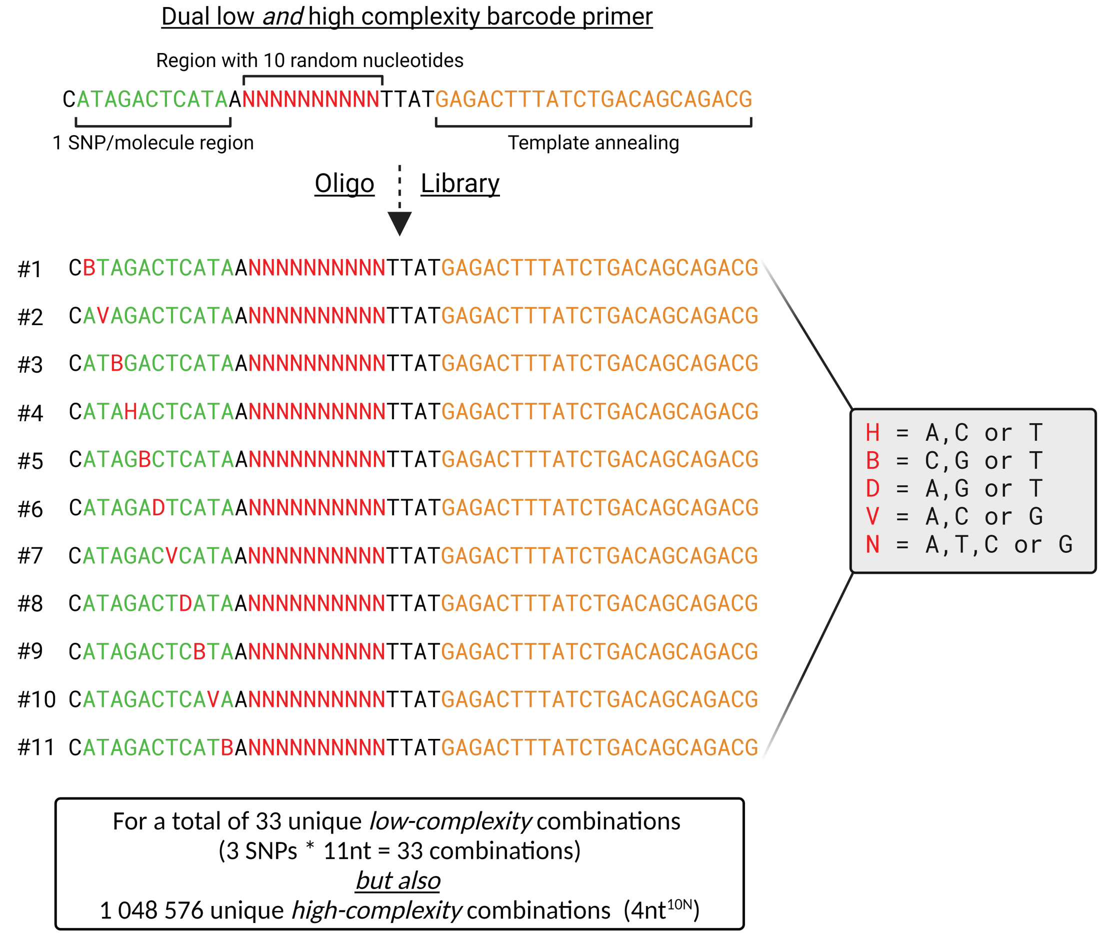


B

| 33 SNPs Barcode | Plasmids/cell | | | | | |
|---|---|---|---|---|---|---|
| | 1 Plasmid | 2 Plasmids | 3 Plasmids | 4 Plasmids | 5 Plasmids | 6 Plasmids |
| Probability that all plasmids have unique SNPs | 1.00 | 0.97 | 0.91 | 0.83 | 0.73 | 0.62 |
| Probability of at least one SNP recurring more than once | 0.00 | 0.03 | 0.09 | 0.17 | 0.27 | 0.38 |

| 10N Barcode | Number of random nucleotides | | | | | | | | | |
|---|---|---|---|---|---|---|---|---|---|---|
| | 1 | 2 | 3 | 4 | 5 | 6 | 7 | 8 | 9 | 10 |
| | Probability that at least 1 random nucleotide is unique | | | | | | | | | |
| 1 Plasmid/cell | 1.00 | 1.00 | 1.00 | 1.00 | 1.00 | 1.00 | 1.00 | 1.00 | 1.00 | 1.00 |
| 2 Plasmids/cell | 0.75 | 0.94 | 0.98 | 1.00 | 1.00 | 1.00 | 1.00 | 1.00 | 1.00 | 1.00 |
| 3 Plasmids/cell | 0.38 | 0.61 | 0.76 | 0.85 | 0.90 | 0.94 | 0.96 | 0.98 | 0.99 | 0.99 |
| 4 Plasmids/cell | 0.09 | 0.18 | 0.26 | 0.33 | 0.39 | 0.45 | 0.50 | 0.55 | 0.59 | 0.63 |

**Figure S1.**

The design of our dual-barcoded library. A) Schematic illustration of the 11 different oligo sequences required to create our dual-barcoded plasmid library. Illustration created with BioRender.com. B) Probabilities of correctly enumerating the number of plasmids per cell if using Sanger sequencing and either the SNP or high-diversity (10N) barcode approach.

Fig. S2

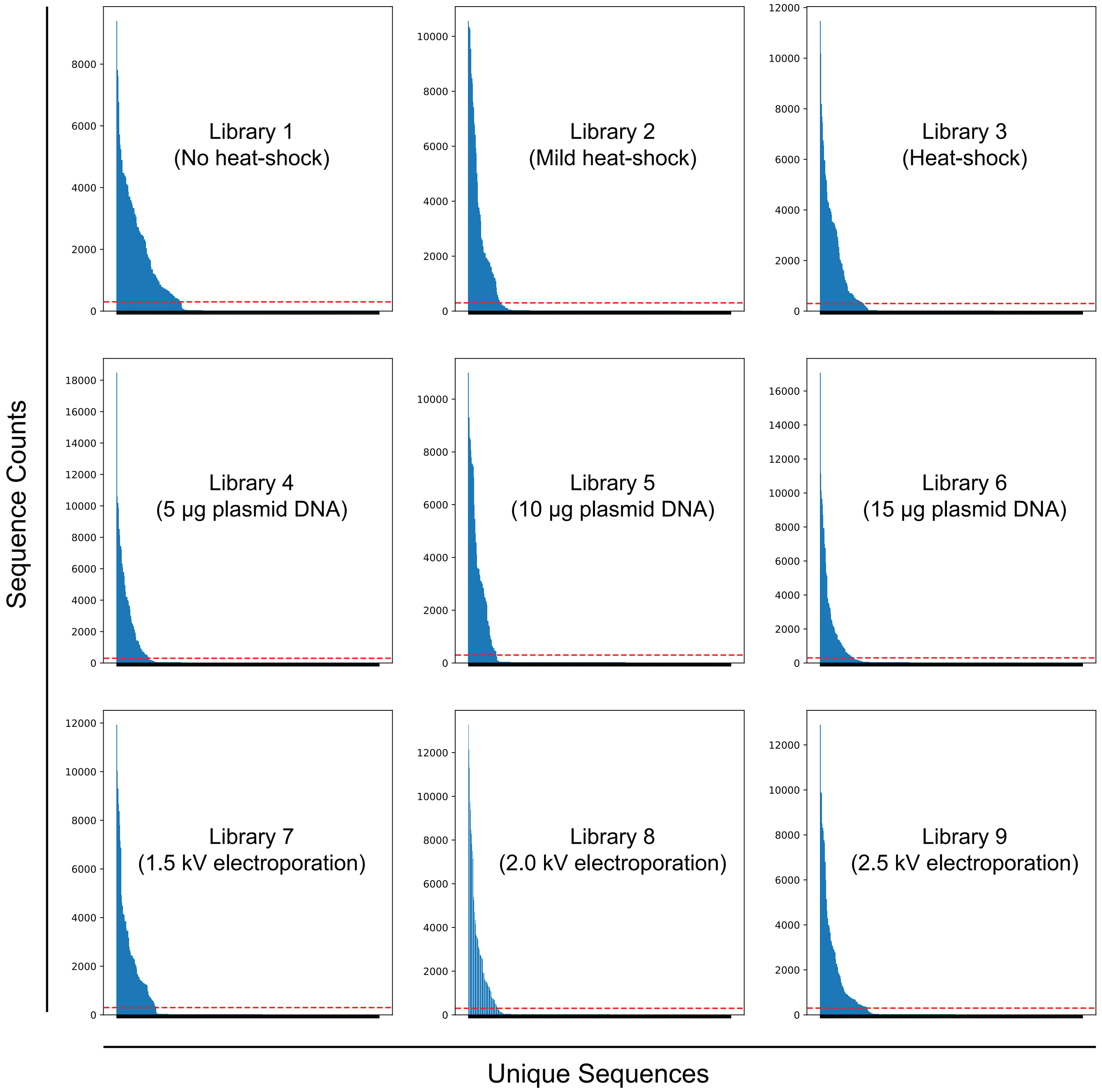
Sequence Counts
Unique Sequences
Library 1
(No heat-shock)
Library 2
(Mild heat-shock)
Library 3
(Heat-shock)
Library 4
(5 µg plasmid DNA)
Library 5
(10 µg plasmid DNA)
Library 6
(15 µg plasmid DNA)
Library 7
(1.5 kV electroporation)
Library 8
(2.0 kV electroporation)
Library 9
(2.5 kV electroporation)

**Figure S2.**

A 300-fold coverage best fits the distribution of the NGS data. The number of identified unique barcodes and the occurrence, i.e., coverage, of each sequence in all nine NGS libraries (experimental conditions). The dashed red lines represent a 300-fold coverage.
